# Glutamate-ammonia ligase promotes lung cancer cell growth through an enzyme-independent upregulation of CaMK2G under a glutamine-sufficient condition

**DOI:** 10.1101/818575

**Authors:** Jiangsha Zhao, Xiankun Zeng, Steven X. Hou

## Abstract

Glutamate-ammonia ligase (GLUL) is highly expressed in many cancer cells. Synthesizing glutamine by its enzyme function has been found to be important for supporting cancer cell survival and growth under glutamine restriction. However, GLUL’s functions under a glutamine-sufficient condition still have not been uncovered. Here we find that GLUL is highly expressed in lung cancer cells and provides survival and growth advantages under both glutamine restriction and adequacy conditions. Knocking down GLUL can block lung cancer cell growth in an enzyme-independent way when glutamine is sufficient. Mechanistically, GLUL regulates Calcium/Calmodulin Dependent Protein Kinase II Gamma (CaMK2G) expression at the transcription level, and CaMK2G is a major mediator in controlling cell growth under GLUL. The transcriptional regulation of CaMK2G is partially mediated by SMAD4. Our data unveil a new enzyme-independent function of GLUL in lung cancer cells under a glutamine-sufficient condition.

## INTRODUCTION

Cancer cells remodel glutamine metabolism to acquire advantages in growth and survival. After intake by cancer cells, glutamine can be directly used for nucleotide and uridine diphosphate-N-acetylglucosamine (UDP-GlcNAc) synthesis, catabolized and used for glutathione synthesis, or used to provide an ATP and carbon source through the tricarboxylic acid (TCA) cycle (Altman et al., 2016). To support their high demand for glutamine, cancer cells incept a large amount of glutamine from the surrounding microenvironment, including stromal tissue (Fuchs and Bode, 2005; Yang et al., 2016). However, cancer cells can also produce small amounts of glutamine through proteolysis and de novo synthesis to maintain survival when extracellular glutamine is limited (Lin et al., 2012; Zhang et al., 2017). The rate-limiting enzyme that catalyzes glutamine synthesis is glutamate-ammonia ligase (GLUL), also known as glutamine synthetase (GS).

GLUL synthesizes glutamine from glutamate and ammonia. It is majorly expressed in normal muscle tissue, adipose tissue, the brain, and the liver to balance ammonia, glutamate, and glutamine levels in those organs and the whole body (Curthoys and Watford, 1995). The expression of GLUL is relatively low in other normal tissues but was found to be elevated in cancer cells from the liver, brain, and breasts (Christa et al., 1994; Rosati et al., 2013; Wang et al., 2016). Since GLUL is the only glutamine synthetase in the human body, most studies focus on its functions in benefiting cancer cells’ survival and growth under glutamine restriction. However, there are many clues that indicate highly expressed GLUL in cancer cells may have extra roles. First, cancer cells maintain a high expression of GLUL even when glutamine is sufficient. The expression of GLUL is tightly controlled. A low level of GLUL can be maintained through glutamine-induced feedback degradation of GLUL under a glutamine-sufficient condition (Arad et al., 1976; Nguyen et al., 2016). However, some cancer cells have a relatively high GLUL expression even when they are supplied with enough glutamine (Bott et al., 2015; Yang et al., 2014). Second, both GLUL and glutamine catabolic enzymes like glutaminase (GLS) are highly expressed in the same cancer cells (Bott et al., 2015; Kung et al., 2011; Tardito et al., 2015). High GLS expression can facilitate cancer cells catalyzing glutamine into glutamate, then glutamate can be used for downstream processes. However, highly expressed GLUL will convert glutamate back into glutamine. An ineffective loop will be formed if both enzymes maintain their catalytic activity when glutamine is sufficient. Third, GLUL’s enzyme function is not necessary for cancer cells when glutamine is sufficient. Although GLUL plays an important role in supporting cell survival and growth under glutamine restriction, blocking its enzyme function by methionine sulfoximine (MSO), a specific GLUL inhibitor has no effect on cell growth and survival when glutamine is sufficient (Tardito et al., 2015; Yang et al., 2014). All these phenotypes indicate that GLUL may have some enzyme-independent functions in cancer cells when glutamine is sufficient.

In this study, we used non-small lung cancer cells as a model and analyzed GLUL’s functions under glutamine restriction or adequacy conditions. We found that GLUL can promote lung cancer cell proliferation through an enzyme-independent increase of CaMK2G expression when glutamine is sufficient. We further found that the regulation of CaMK2G by GLUL is partially mediated by SMAD4. Our results uncover a previously unreported enzyme-independent function of GLUL under a glutamine-sufficient condition.

## RESULTS

### GLUL is highly expressed in lung cancer cells

To analyze the expression of GLUL in lung cancer samples, a human lung cancer tissue microarray was checked by immunohistochemistry. The GLUL expression was found to be low in normal lung tissues (Figure 1A upper left and 1B) but much higher in all kinds of lung cancer samples (Figure 1A and 1B). We also noticed that there is a heterogenous expression of GLUL in many cancer samples, in which GLUL-positive cancer cells are surrounded by huge numbers of GLUL-negative cancer cells (Figure S1). As the clinical stage information was also available, we further analyzed the correlation between the GLUL level and the clinical stage of those samples. Although there was no significant correlation in small cell undifferentiated carcinoma (SCC), a positive correlation could be found in non-small-cell carcinoma (NSCC) samples (Figure 1C and 1D). We then chose to focus on NSCC and further checked GLUL expression in eight established non-small lung cancer cell lines. As detected by Western blotting, five lung cancer cell lines showed a high GLUL expression (Figure 1E). As we could detect GLUL-positive cancer cells in those GLUL-low samples, we wanted to know whether those GLUL-low and GLUL-negative lung cancer cell lines also contained few GLUL-positive cells. We then isolated single-cell clones from A549, a cell line with an extremely low level of GLUL, and detected their GLUL expression. Interestingly, we found few clones with a relatively high expression of GLUL, while all of the remaining clones, as well as the parental A549 cells, were GLUL-negative (Figure 1F).

**Figure 1.**
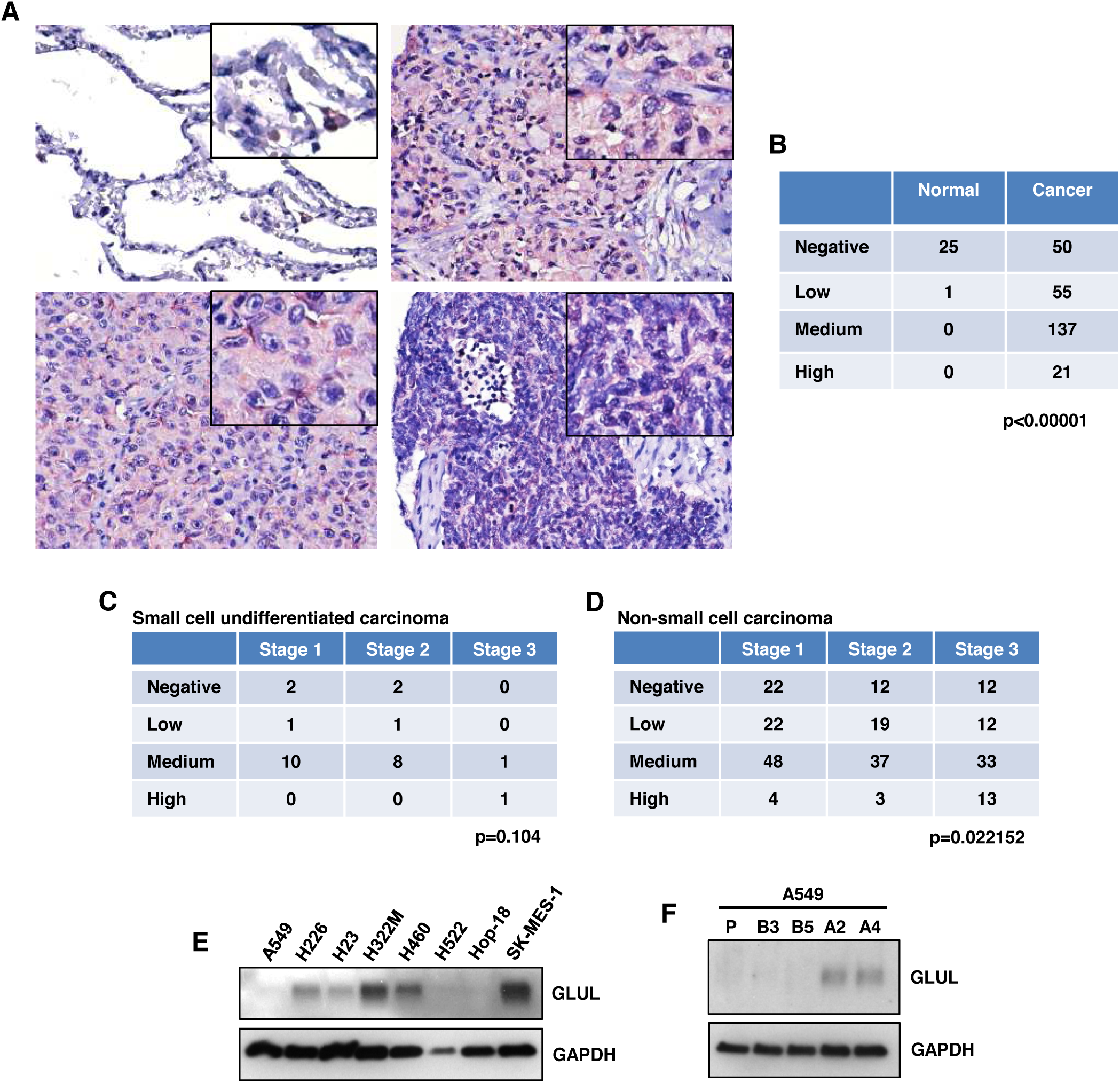
GLUL is highly expressed in lung cancer cells. (A) Representative IHC staining of GLUL in normal lung tissue and lung cancer tissues. Upper left: normal lung; upper right: adenocarcinoma; bottom left: squamous cell carcinoma; bottom right: small cell undifferentiated carcinoma. Inserts: enlarged pictures to show GLUL staining. Scale bar: 50 micrometers. (B) Summarized GLUL expression data in clinical samples. (C) The co-relationship between GLUL expression and the clinical stage in small cell undifferentiated carcinoma. (D) The co-relationship between GLUL expression and the clinical stage in non-small cell carcinoma. (E) GLUL protein expression in non-small lung cancer cell lines. (F) GLUL protein expression in A549 parental and single-cell derived clones.

### GLUL is not necessary for cell survival after glutamine deprivation in some GLUL-positive lung cancer cells

GLUL has been reported to be important for sustaining proliferation and survival after glutamine starvation in other cancer types (Bott et al., 2015; Kung et al., 2011; Tardito et al., 2015). Consequently, we checked whether this would be the same in lung cancer cells. Among the six lung cancer cell lines, the GLUL-negative A549 parental line and B5 clone showed a dramatic cell number reduction and a dead cell increase after glutamine deprivation (Figure 2A and 2B). However, no such dramatic change of cell number and cell death was detected in the other four GLUL-positive lung cancer cell lines (Figure 2A and 2B). Although a slight increase in cell number was detected in H322M after glutamine deprivation, the cell number of the other three GLUL-positive cell lines did not increase the after glutamine deprivation. These results suggest that glutamine is necessary for lung cancer cell growth that disregards the GLUL expression level. The resistance of cell death induced by glutamine starvation in GLUL-positive cells further prompted us to ask whether GLUL is necessary for their survival. We then treated the four GLUL-positive lung cancer cells with MSO, a specific GLUL inhibitor, under glutamine deprivation. Surprisingly, blocking the enzyme function of GLUL with MSO only increased cell death in the A549-A4 and H460 cells but not in SK-MES-1 (Figure 2C). The MSO treatment even slightly reduced cell death in H322M cells (Figure 2C). To further confirm the function of GLUL in cell death resistance under glutamine starvation, we knocked down the GLUL expression in all four GLUL-positive cell lines by using two independent shRNA sequences (Figure 3A). Constantly, we found a significant increase in cell death in the GLUL-knocked-down A549-A4 and H460 cells after glutamine deprivation (Figure 2D). Knocking down GLUL in SK-MES-1 did not affect cell survival, and in H322M cells it also slightly reduced cell death after glutamine deprivation (Figure 2D). These results indicate that GLUL is not necessary for cell survival under glutamine starvation in some GLUL-positive lung cancer cells.

**Figure 2.**
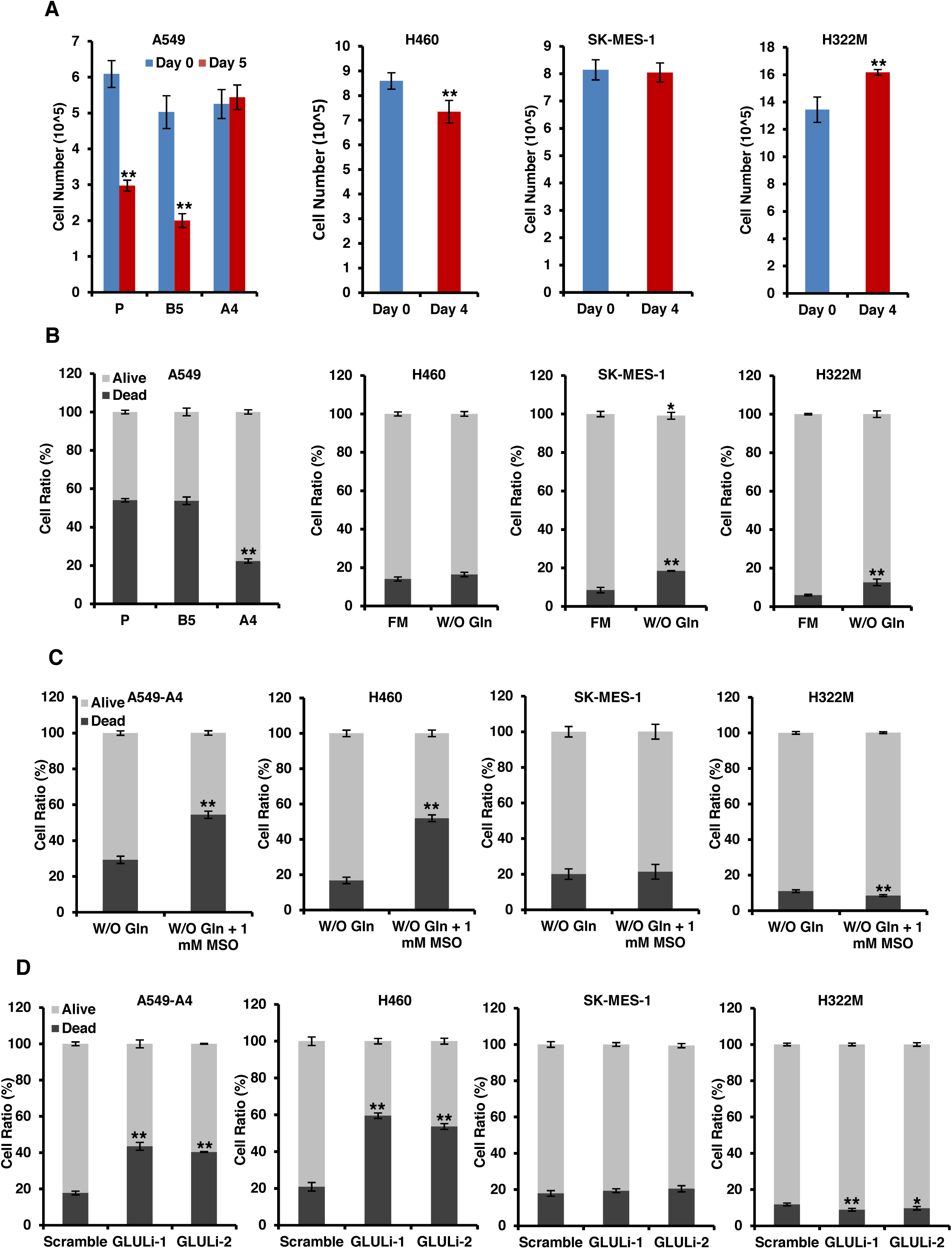
GLUL is not necessary for survival of some GLUL-positive lung cancer cells after glutamine deprivation. (A) Cell growth of lung cancer cells after glutamine deprivation. (B) Cell death of lung cancer cells after glutamine deprivation (same days of treatment as panel (A)). (C) Cell death of lung cancer cells after glutamine deprivation with 1 mM MSO treatment for four days. (D) Cell death of lung cancer cells with GLUL knocking down after glutamine deprivation for four days. *p < 0.05, **p < 0.01.

**Figure 3.**
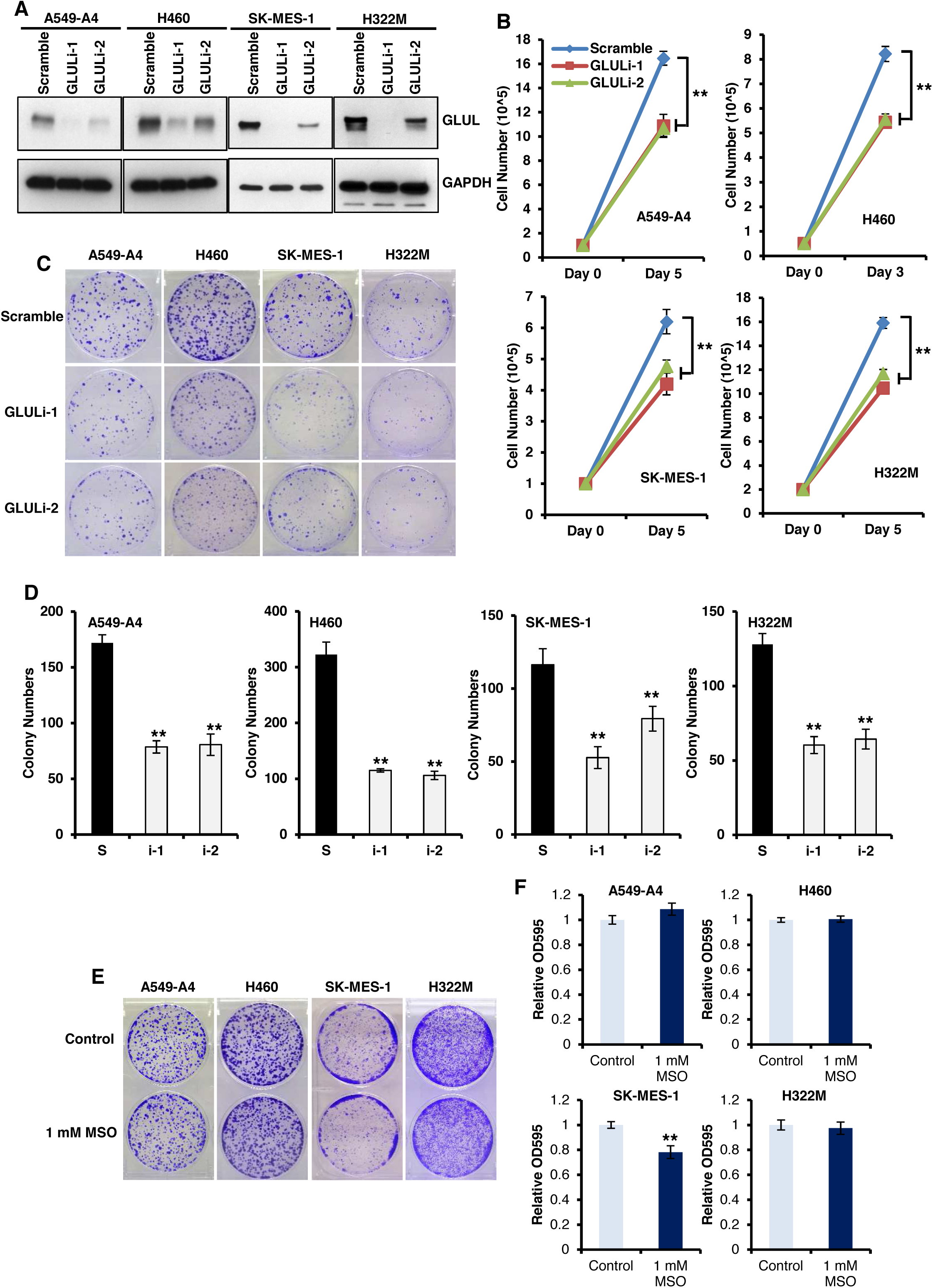
Knocking down GLUL blocks cell growth in lung cancer cells under an FM condition. (A) GLUL protein expression in GLUL-knocked-down lung cancer cells. (B) Cell growth of scramble cells and GLUL-knocked-down lung cancer cells under an FM condition. The same number of cells were seeded at day 0, and the cell number was counted after 3–5 days’ culture under an FM condition. (C) Representative crystal violet staining of colonies from scramble cells and GLUL-knocked-down lung cancer cells after 10– 14 days’ culture under an FM condition. (D) Quantification of panel (C) by counting the colony’s number. (E) Representative crystal violet staining of colonies from control and 1 mM MSO–treated lung cancer cells under an FM condition for 10–14 days. (F) Quantification of panel (E) by OD595 absorption. *p < 0.05, **P < 0.01.

### GLUL-promoted cancer cell growth under a glutamine-sufficient condition is independent of its enzyme function

When we cultured the cells in full medium (FM), we noticed that the GLUL-knocked-down cells had a more diminished cell growth than the scrambled control. As the FM contains intact glutamine supplementation, we asked whether GLUL plays some roles in the cells without glutamine restriction. Interestingly, knocking down GLUL can significantly reduce cell growth under FM, as measured by cell number change in all four lung cancer cell lines we checked (Figure 3B). To further confirm this phenotype, we used a colony formation assay to detect colony growth in those cells. Consistent with the cell number counting assay, we found a dramatic reduction in the colony number of the GLUL-knocked-down lung cancer cells under an FM condition (Figure 3C and 3D). To analyze the mechanism that causes the inhibition of cell growth in GLUL-knocked-down cells under an FM condition, we checked the cell death and cell cycle in those cells. There is a notable decrease in the G0/G1 phase ratio but no change of cell death in the GLUL-knocked-down cells (Figure S2A and S2B). As FM contains sufficient glutamine for cancer cells, it is interesting to explore whether the enzyme function of GLUL is needed for its growth promotion effect in lung cancer cells. Surprisingly, blocking the enzyme function of GLUL in lung cancer cells by using MSO did not reduce cell growth under an FM condition in the A549-A4, H460, and H322M cells (Figure 3E and 3F). The only exception is the SK-MRES-1 cells, in which MSO treatment slightly reduced the cell growth. These results suggest that GLUL can play an enzyme-independent role in regulating cell growth under an FM condition.

### GLUL actives CaMK2G to promote cell growth under a glutamine-sufficient condition

To investigate the potential molecular mechanism that may regulate the enzyme-independent function of GLUL in cell growth, we checked some signaling pathways that have been reported to regulate the cell growth of lung cancer cells. Although we did not find a notable change in the AKT, mTOR, and ERK pathways (Figure S3), we detected a dramatic reduction of phosphorylated Ca2+/calmodulin-dependent kinase II (CaMK2) in the GLUL-knocked-down cells cultured with FM (Figure 4A). CaMK2 is a multifunctional serine/threonine kinase (Braun and Schulman, 1995). There are four major isotypes of CaMK2: CaMK2A, CaMK2B, CaMK2D and CaMK2G. To identify which CaMK2 isotype has been affected by GLUL in lung cancer cells, we detected their mRNA expression in the lung cancer cells and found that CaMK2G is the major isoform expressed in the lung cancer cell lines (Figure 4B). We then checked CaMK2G protein expression in the GLUL-knocked-down cells. Consistent with the reduction of phosphorylated CaMK2, the total level of CaMK2G was also dramatically reduced in the GLUL-knocked-down cells (Figure 4A). We then treated the lung cancer cells with a specific CaMK2 inhibitor, KN93, to mimic the blockage of CaMK2G activity in the GLUL-knocked-down cells. When compared to the non-effect control compound KN92, KN93 treatment can significantly reduce cell growth in lung cancer cells (Figure 4C). To further confirm the function of GLUL in regulating cell growth is through CaMK2G activation, we forced an overexpression of CaMK2G in the SK-MES-1 scramble and GLUL-knocked-down cells. The overexpression of CaMK2G dramatically increased the phosphorylated CaMK2G in all three cells (Figure 4D) and overcame the colony number reduction caused by knocking down GLUL (Figure 4E and 4F). It is already reported that CaMK2G is highly expressed in NSCC cells and promotes their survival and growth (Chai et al., 2014; Chai et al., 2015). Our results indicate that CaMK2G is a target of GLUL in regulating lung cancer cell growth under a glutamine-sufficient condition.

**Figure 4.**
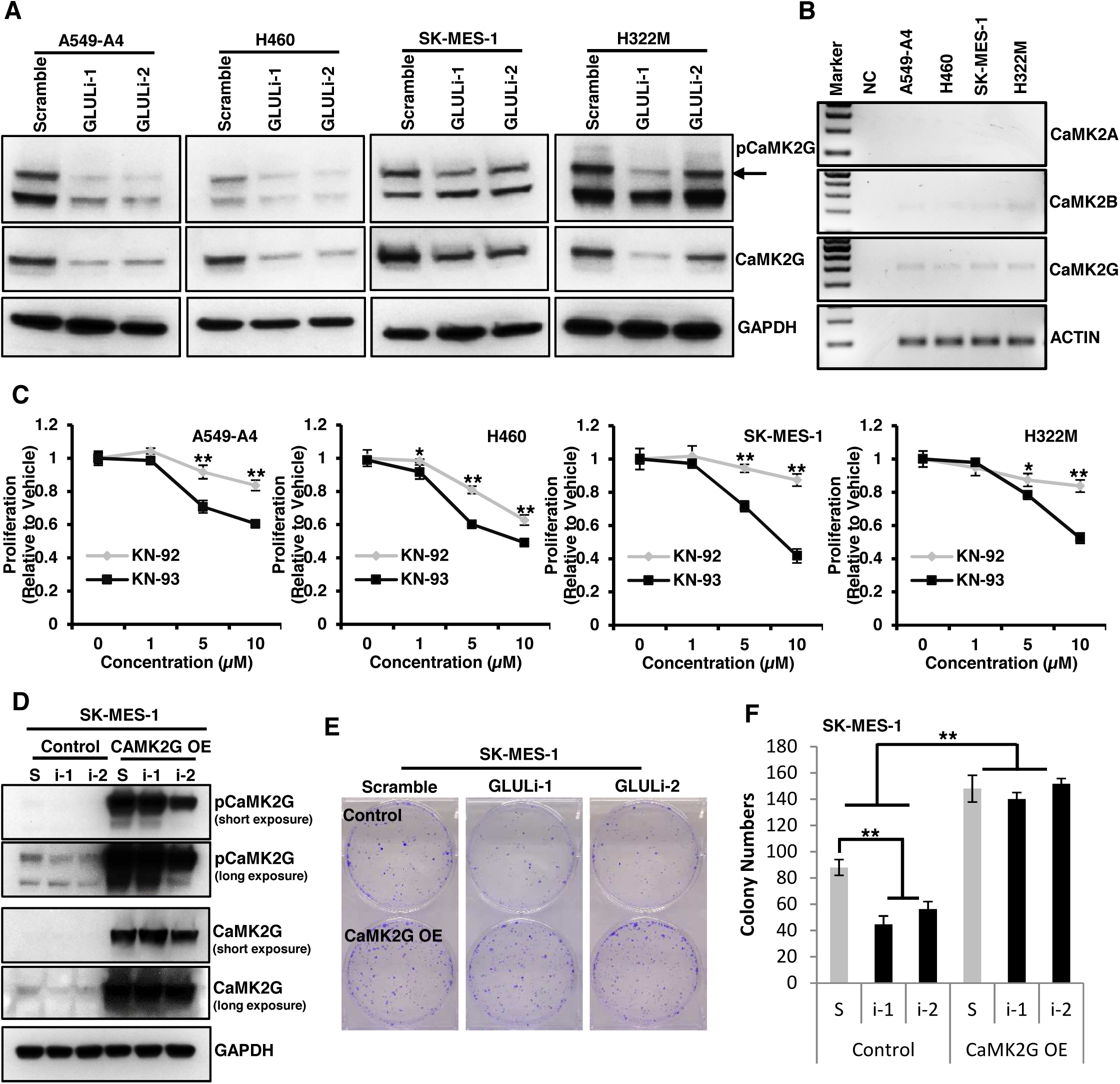
CaMK2G is a downstream target of GLUL. (A) Phosphorylated and total CaMK2G protein expression in GLUL-knocked-down lung cancer cells. (B) mRNA level of CaMK2 isoforms in lung cancer cells was detected by PCR. (C) Cell growth of lung cancer cells treated with KN-92 and KN-93 for two days. (D) Phosphorylated and total CaMK2G protein expression in control and CaMK2G-overexpressed (OE) cells. (E) Representative crystal violet staining of colonies from control and CaMK2G-overexpressed (OE) cells. (F) Quantification of panel (E) by counting the colony’s number. *p < 0.05, **p < 0.01.

### GLUL regulates CaMK2G expression at the transcription level

The reduction of the CaMK2G total protein level in the GLUL-knocked-down cells raised the question of how GLUL modifies CaMK2G expression. Gene expression can be regulated both in mRNA and at the protein level. At the protein level, protein degradation through proteasomes or lysosomes is a major way to control protein’s abundance in cells. We first checked whether downregulation of CaMK2G in the GLUL-knocked-down cells was through protein degradation. We treated the cells with MG132 or chloroquine to block protein degradation by proteasomes or lysosomes, respectively. No rescue of the CaMK2G protein level could be detected after either MG132 or chloroquine treatment in the GLUL-knocked-down cells (Figure 5A and 5B). These results indicate that the downregulation of CaMK2G in the GLUL-knocked-down cells is not through protein degradation pathways. We then checked the expression of CaMK2G at the transcription level. Indeed, we found a significant reduction of CaMK2G mRNA in the GLUL-knocked-down cells (Figure 5C).

**Figure 5.**
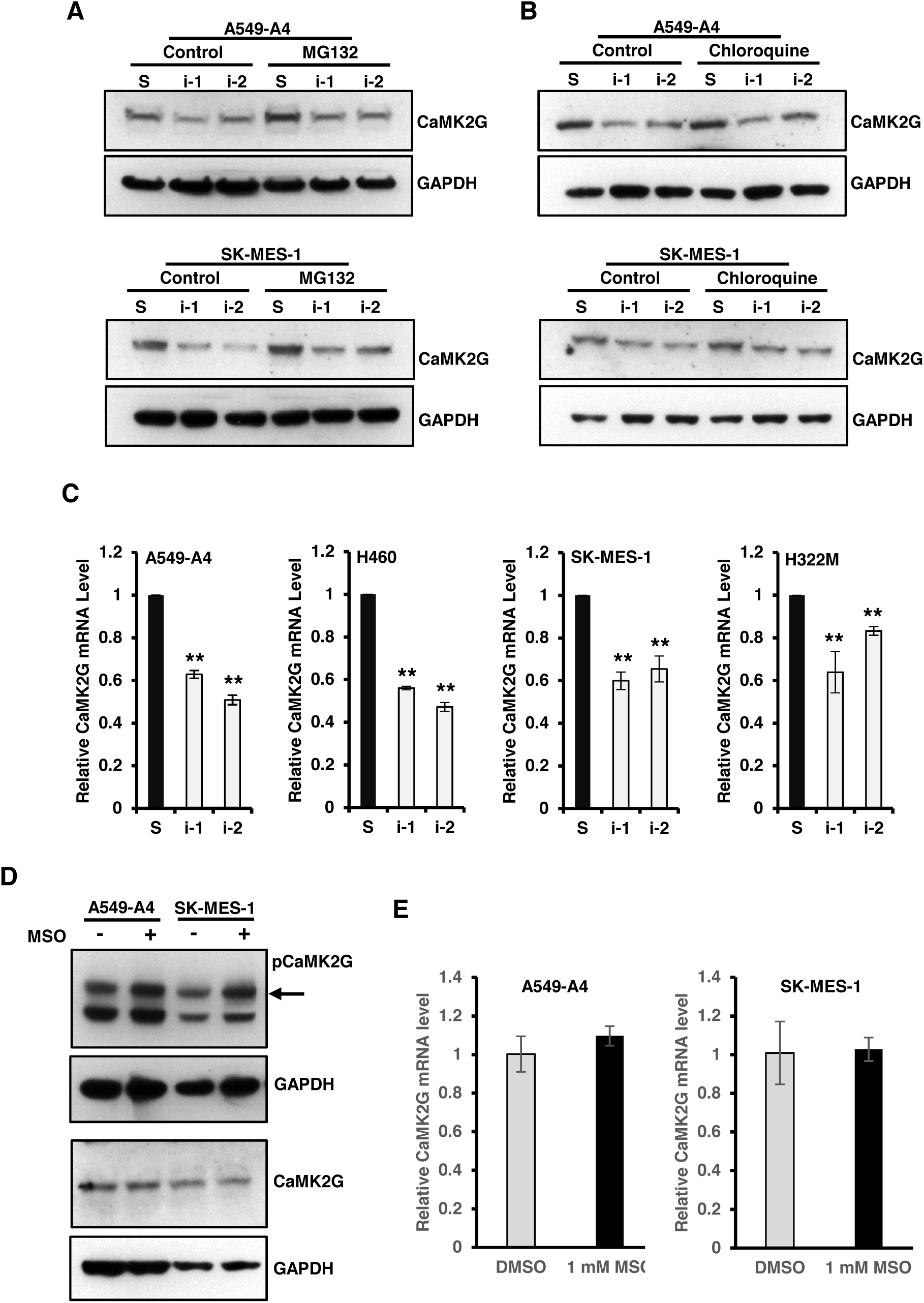
GLUL regulates the transcription of CaMK2G in an enzyme-function-independent manner. (A) Total CaMK2G protein expression in GLUL-knocked-down lung cancer cells treated with dimethyl sulfoxide (DMSO) or 1 µg/ml MG132 for one day. (B) Total CaMK2G protein expression in GLUL-knocked-down lung cancer cells treated with DMSO or 100 µM chloroquine for one day. (C) mRNA level of CaMK2G in GLUL-knocked-down lung cancer cells (real-time PCR). (D) Phosphorylated and total CaMK2G protein expression in control and 1 mM MSO–treated cells. (E) mRNA level of CaMK2G in control and 1 mM MSO–treated cells (real-time PCR). **p < 0.01.

As we already found that GLUL promotes the growth of lung cancer cells under a glutamine-sufficient condition independently of its enzyme function, we asked whether the activation of CaMK2G by GLUL under FM also occurs through the same way. As expected, MSO treatment did not reduce, but slightly increased, the phosphorylated CaMK2G with no change in its total level (Figure 5D). Consistent with this MSO treatment also did not change the mRNA level of CaMK2G in lung cancer cells (Figure 5E).

### GLUL controls the translocation of SMAD4

The reduced CaMK2G mRNA level indicates GLUL can regulate its transcription through direct or indirect ways. As GLUL majorly localizes in cytosol and because we did not find a notable level of GLUL in the nucleus (data not shown), there must be a transcription factor (TF) involved in regulating CaMK2G’s transcription by GLUL. To identify the potential TFs that can be regulated by GLUL, we carried out a TF activation profiling array in the SK-MES-1 scramble and GLUL-knocked-down (shRNA 1#; i-1) cells. As shown in Figure 6A and 6B, five out of forty-seven checked TFs showed a notable activity reduction (>30%). However, only two of them, Pit-1 and SMAD, reached the cutoff line (two-fold change) suggested by the manufacturer. Pit-1 (also known as POU1F1) is a pituitary-specific transcription factor. It is majorly expressed in the pituitary gland and regulates pituitary development and hormone expression (Simmons et al., 1990). Consequently, we chose to check SMAD further.

**Figure 6.**
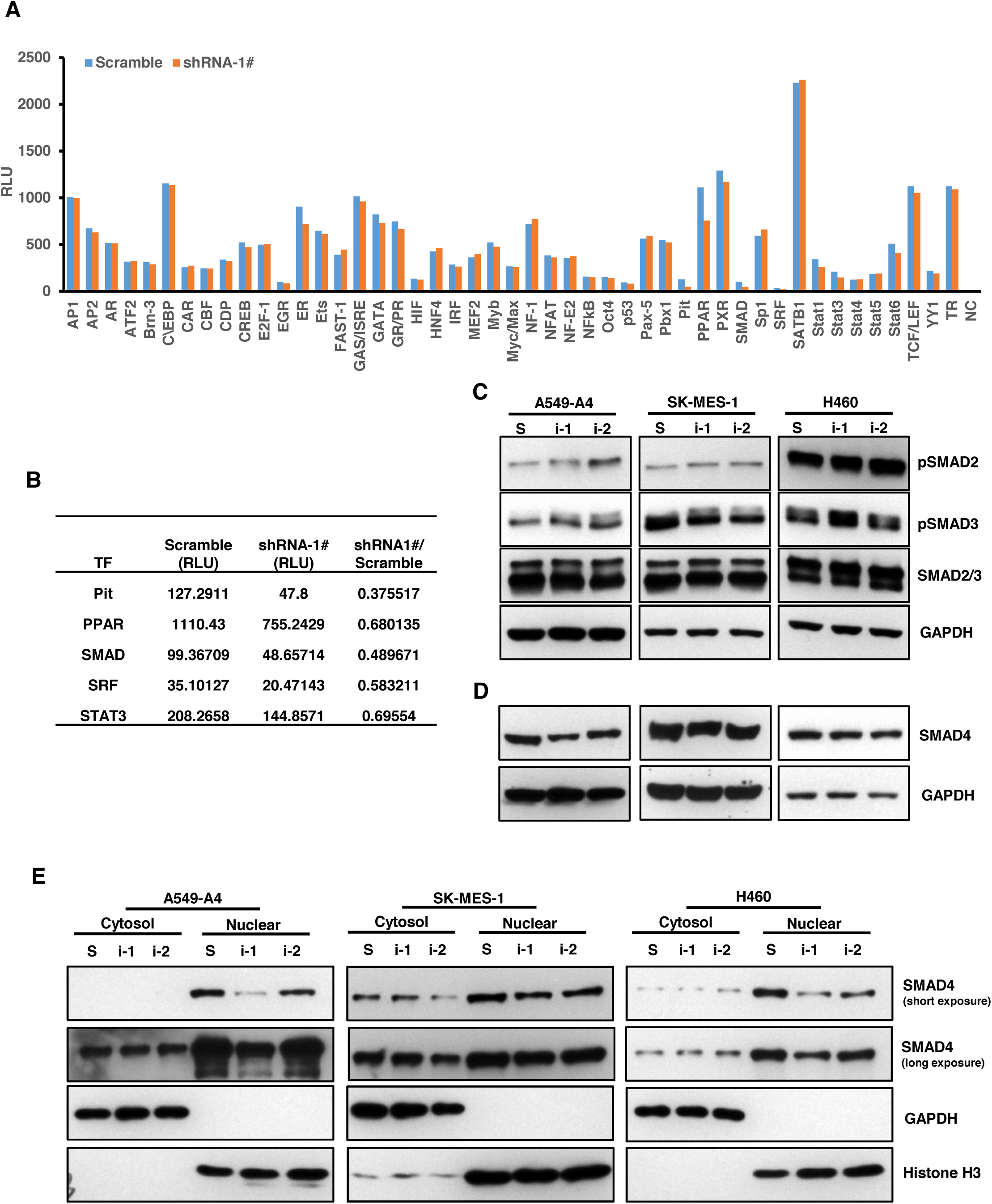
GLUL regulates the expression and translocation of SMAD4. (A) The change of TF activity in GLUL-knocked-down SK-MES-1 cells. Forty-seven TFs were measured by TF activation profiling array. (B) The top five changed TFs detected in panel (A). (C) Phosphorylated and total SMAD2/3 protein expression in GLUL-knocked-down lung cancer cells. (D) Total SMAD4 protein expression in GLUL-knocked-down lung cancer cells. (E) Cytoplasmic and nuclear SMAD4 protein expression in GLUL-knocked-down lung cancer cells.

SMAD is a large protein family that contains eight members: the receptor-regulated SMADs (R-SMADs) SMAD1, SMAD2, SMAD3, SMAD5, and SMAD8 (SMAD9); the common mediator SMAD (co-SMAD) SMAD4; and the inhibitory SMADs (I-SMADs) SMAD6 and SMAD7 (Schmierer and Hill, 2007). As the TF activation profiling array cannot distinguish the individual SMAD proteins, we should further identify the actual target of GLUL in lung cancer cells. The transforming growth factor beta (TGF-beta) pathway is one of the major pathways involved in cancer initiation and progression, including lung cancer (Massague, 2008). We then chose to test SMAD2, SMAD3 and SMAD4 the downstream mediators of the TGF-beta pathway, first. Activation of the TGF-beta pathway will phosphorylate SMAD2 and SMAD3, and the phosphorylated SMAD2 and SMAD3 then bond with SMAD4 and translocate into the nucleus to be activated as a transcriptional complex. To analyze the activation of SMAD2 and SMAD3, we checked their phosphorylation level in the GLUL-knocked-down cells, and no notable change has been found (Figure 6C). Furthermore, no difference was found in total SMAD2/3 proteins (Figure 6C). However, we found a notable reduction in SMAD4 proteins in the GLUL-knocked-down cells (Figure 6D). As the only co-SMAD, SMAD4 needs to be translocated into the nucleus to accomplish its transcriptional functions. We then checked whether the translocation of SMAD4 was also affected by GLUL. Indeed, we found a dramatic reduction in the nuclear SMAD4 protein level in the GLUL-knocked-down cells (Figure 6E). Interestingly, we did not find any notable change in the cytoplasmic SMAD4 level in the GLUL-knocked-down cells while the total and nuclear SMAD4 proteins were reduced (Figure 6F).

We then checked whether the regulation of SMAD4 by GLUL is dependent on its enzyme function. We found that blocking GLUL’s enzyme function by MSO did not change the SMAD4 protein level in lung cancer cells (Figure 7A). These results indicate that GLUL can regulate the total SMAD4 level and its location in lung cancer cells in an enzyme-independent way.

**Figure 7.**
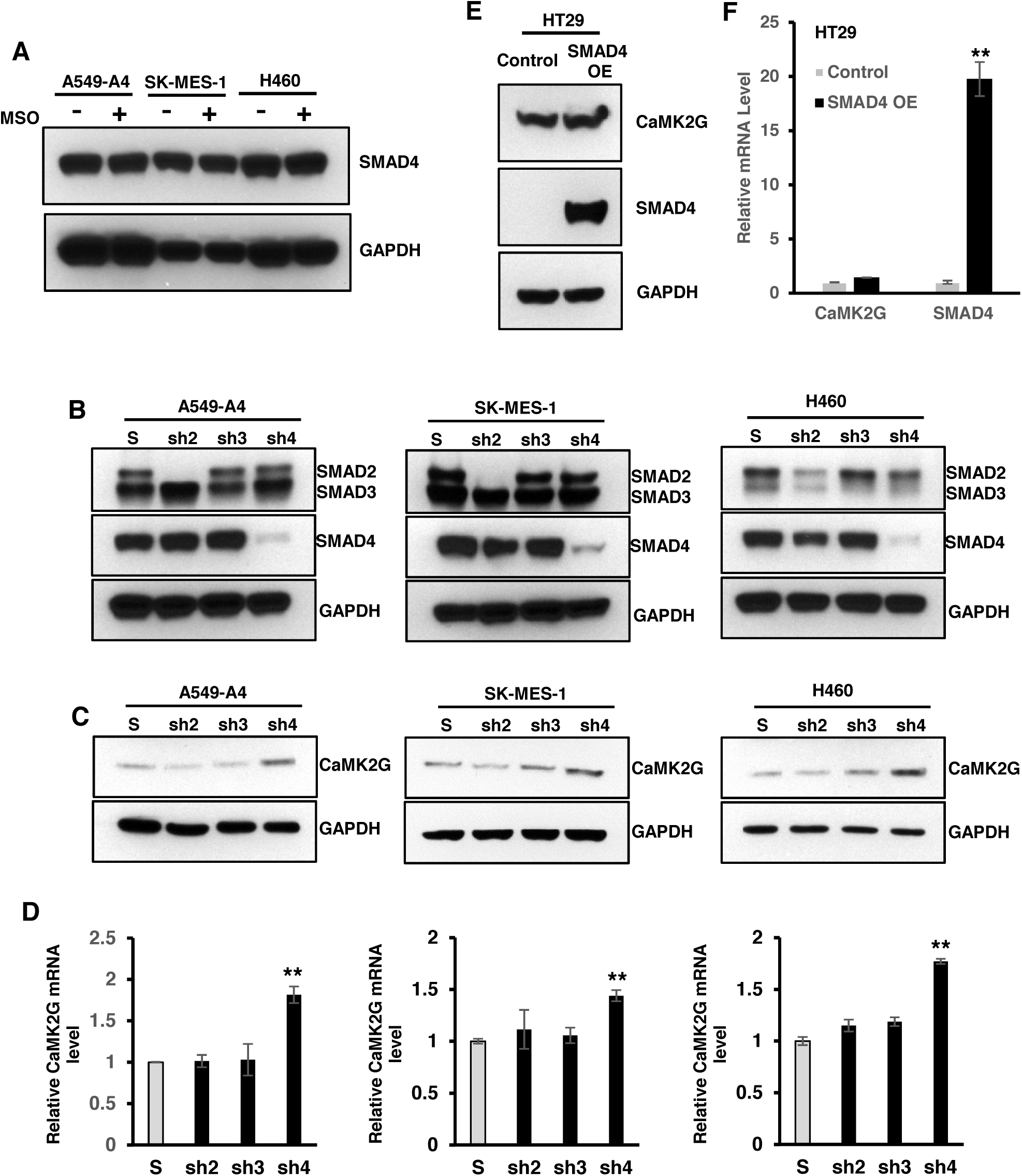
SMAD4 regulates the expression of CaMK2G at the transcription level. (A) Total SMAD4 protein expression in control and 1 mM MSO–treated cells. (B) Total SMAD2/3/4 protein expression in SMAD2-, SMAD3-, or SMAD4-knocked-down lung cancer cells. (C) Total CaMK2G protein expression in SMAD2-, SMAD3-, or SMAD4-knocked-down lung cancer cells. (D) mRNA levels of CaMK2G in SMAD2-, SMAD3-, or SMAD4-knocked-down lung cancer cells (real-time PCR). (E) Total CaMK2G and SMAD4 protein expression in control and SMAD4-overexpressed (SMAD4 OE) HT29 cells. (F) mRNA levels of CaMK2G and SMAD4 in control and SMAD4-overexpressed HT29 cells. S: scramble; sh2: shSMAD2; sh3: shSMAD3; sh4: shSMAD4. **p < 0.01.

### SMAD4 regulates CaMK2G transcription

To analyze whether SMAD4 is involved in regulating CaMK2G transcription, we knocked down SMAD2, SMAD3, and SMAD4 in lung cancer cells and checked their effects in CaMK2G expression. All constructs reduced their targeting proteins dramatically (Figure 7B). However, knocking down SMAD2 or SMAD3 had no effect on CaMK2G expression in protein or mRNA levels (Figure 7C and 7D). These results further confirm that SMAD2/3 are not the downstream targets of GLUL. Surprisingly, we found a significant increase of CaMK2G mRNA and protein levels in SMAD4-knocked-down cells (Figure 7C and 7D). Although these results indicate that SMAD4 is involved in regulating CaMK2G, they were the opposite of what we expected. Knocking down SMAD4 by shRNA is supposed to reduce SMAD4 protein in both the cytoplasm and nucleus. Consistent with this, we found that knocking down GLUL reduces SMAD4 protein in the nucleus but not in the cytoplasm. As we found an opposite effect in CaMK2G’s transcription by knocking down GLUL and SMAD4, we hypothesized that the cytoplasmic, but not nuclear, SMAD4 is important for the regulation of CaMK2G. In other words, SMAD4 should not be a direct factor that regulates the transcription of CaMK2G. Consistent with this hypothesis, we found that overexpressing SMAD4 in SMAD4-null colon cancer cell line HT29 did not show dramatic change in CaMK2G expression (Figure 7E and 7F). These results indicate that SMAD4 is involved in regulating CaMK2G transcription by GLUL through an indirect mechanism.

## DISCUSSION

Highly expressed GLUL has been found in many types of cancer cells, and it plays important roles in maintaining cell survival and growth under glutamine deprivation (Bott et al., 2015; Cox et al., 2016; Tardito et al., 2015). However, the functions of GLUL in cancer cells with sufficient glutamine are still unrevealed. In this study, we have found an enzyme-independent function of GLUL in regulating lung cancer cell proliferation when sufficient glutamine is supplied. Our findings expand the knowledge about GLUL’s functions in cancer cells. First, our results show that GLUL can regulate cell survival and growth not just under glutamine restriction, but also under a glutamine adequacy condition. Second, GLUL plays its roles in both enzyme-dependent and independent ways. Third, our results indicate that targeting GLUL to degradation will be more efficient than just targeting its enzyme function during cancer therapy.

GLUL expression is tightly controlled in bacteria through different feedforward and feedback mechanisms (Stadtman, 2001). Although the mechanisms regulating GLUL expression in mammalian cells are not as clear as in prokaryotes, a growing body of evidence suggests there is a similar controlling network in mammalian cells. For example, glutamine can work as a major feedback repressor in both bacterial and mammalian cells (Crook and Tomkins, 1978; Streicher and Tyler, 1981). In mammalian cells, high extracellular glutamine can induce degradation of GLUL, which in turn keeps a low GLUL expression level when glutamine is sufficient (Nguyen et al., 2016). As the human body has a relatively high concentration of free glutamine, it is unsurprising that most human organs have a relatively low GLUL expression. In human lung tissue, the glutamine flux is balanced under normal conditions, but a significantly elevated glutamine release can be detected under stress (Hulsewe et al., 2003; Plumley et al., 1990). Although the efflux of glutamine is thought to occur through enhanced proteolysis or de novo synthesis, there is no direct evidence to show the GLUL expression in both normal and stressed human lung tissues. In this study, we have shown that GLUL has a relatively low expression in normal lung tissues. Meanwhile, we found an increased GLUL expression in all subtypes of lung cancer. Elevated GLUL has already been found in many other types of cancers (Christa et al., 1994; Rosati et al., 2013; Wang et al., 2016). Our results indicate that GLUL may also play some important roles in lung cancer progression.

GLUL expression under a glutamine deprivation condition has been found to be important in supporting the survival and growth of many cancer cells. Our results indicate that GLUL is not necessary for the survival of some GLUL-positive lung cancer cells when glutamine is limited. When blocking GLUL’s enzyme function with specific inhibitor MSO or knocking down its expression in four GLUL-positive cell lines, only two lines show significant increases in cell death after glutamine deprivation, and there is no change in the other two cell lines. These results suggest that GLUL’s function is limited under glutamine deprivation in those lung cancer cells. In fact, cancer cells can utilize many other ways to sustain cell proliferation and survival when deprived of glutamine (Cheng et al., 2011; Reid et al., 2016).

Although GLUL’s expression can be repressed by glutamine, we and other researchers found a relatively high level of GLUL in cancer cells even when glutamine is sufficient (Arad et al., 1976; Nguyen et al., 2016). This may be due to the activation of GLUL’s transcription by the enhanced oncogenic pathways like beta-Catenin, AKT, Myc, and GNC2 (Bott et al., 2015; Cadoret et al., 2002; Kung et al., 2011; van der Vos et al., 2012; Yuneva et al., 2012; Zucman-Rossi et al., 2007). As a glutamine synthetase, GLUL’s functions in cancer cells under a glutamine deprivation condition are majorly studied. However, little is known about its functions under a glutamine-sufficient condition. In this study, we reported a new function of GLUL in lung cancer cells with sufficient glutamine. Our data show that GLUL can promote lung cancer cell growth by regulating CaMK2G transcription in an enzyme-function-independent way. These results explain why cancer cells keep a high level of GLUL even when glutamine is sufficient. Our data, in combination with previous findings, clearly show that GLUL can mediate cancer cell survival and growth under both glutamine restriction and sufficiency through enzyme-dependent or independent ways. These functions of GLUL may be extremely important for in vivo cancer tissues. As cancer tissues always have large volumes in vivo, the supplies of glutamine should not be equal. Although the glutamine concentration is not reduced in most of the cancer tissues (Fan et al., 2009), a low intratumoral glutamine level has been detected in tumor core regions (Pan et al., 2016; Reid et al., 2013). In those low glutamine core regions, highly expressed GLUL in cancer cells can provide survival and growth advantages through its enzyme-dependent functions. Meanwhile, it can also promote cancer cell growth in surrounding regions where the glutamine is not restricted through its enzyme-independent functions. These dual functions make GLUL an attractive target for cancer therapy and indicate that targeting GLUL to degradation will be more efficient than just targeting its enzyme function.

As a cytoplasmic enzyme, it is still unclear how GLUL regulates CaMK2G’s transcription. Although we have found that SMAD4 should be one of the mediators involved, we still don’t know the detailed mechanism by which it mediates the regulation of CaMK2G’s expression by GLUL. It is impossible that SMAD4 works as a direct regulator here. The first reason is expression of SMAD4 in SMAD4-null HT29 cells did not change the expression of CaMK2G. Second, downregulation of SMAD4 in GLUL- and SMAD4-knocked-down lung cancer cells had the opposite effect of CaMK2G expression. However, we did not find a notable change in the cytoplasmic SMAD4 level in GLUL-knocked-down cells while the total and nuclear SMAD4 proteins were reduced. SMAD4 can constitutively shuttle between the nucleus and the cytoplasm even without TGF-beta activation (Pierreux et al., 2000). The reduction of total and nuclear SMAD4 without change in the cytoplasm indicates a preferred cytoplasmic location of SMAD4 after GLUL knocking down in lung cancer cells. This can be regulated by either enhancing nuclear export or reducing nuclear import. Based on these results, we hypothesize that the translocation of SMAD4 between the nucleus and the cytoplasm is more important in regulating CaMK2G expression than the nuclear SMAD4. As SMAD4 is not the direct mediator for regulating CaMK2G expression, we believe that other transcription factors should be involved in this process. Besides those R-SMADs, SMAD4 can also interact with other TGF-beta pathway–related or unrelated transcription factors like SNON, lymphoid enhancer-binding factor 1/T cell-specific factor (LEF/TCF), etc. (Lim and Hoffmann, 2006; Stroschein et al., 1999). SMAD4 can regulate their translocation and activation. Further studies are still needed to find out which transcription factor is regulated by SMAD4 in GLUL-knocked-down lung cancer cells and how the transcription factor regulates the transaction of CaMK2G.

## AUTHOR CONTRIBUTIONS

J.Z. and S.H. designed the research. J.Z. performed most of the experiments and data analysis. X.Z. performed the IHC staining for clinical samples. J.Z. and S.H. wrote the manuscript.

## ACKNOWLEDGMENTS

This research was supported by the Intramural Research Program of the National Institutes of Health, National Cancer Institute (to SXH). The authors have no potential conflict of interest to disclose. The content of this publication does not necessarily reflect the views or policies of the US Department of Defense or the US Department of the Army.

## Materials and Methods

### Cell lines and single-cell clone isolation

Human lung cancer cell lines A549, H226, H23, H322M, H460, H522, Hop-18, SK-MES-1, and colon cancer cell line HT29 were kindly provided by the Division of Cancer Treatment and Diagnosis Tumor Repository of the National Cancer Institute (NCI) at Frederick, MD. Phoenix ampho and the 293T cell lines were gifts from Dr. P. Charles Lin (NCI at Frederick). All cell lines were cultured in RPMI1640 supplemented with 10% fetal bovine serum and 100 units/ml penicillin/streptomycin at 37°C in a humidified atmosphere containing 5% CO_2_.

Single-cell clones were isolated from A549 cells as previously reported (Li et al., 2008). Briefly, cells were suspended in full medium at ~5 cells/ml, 200 µl medium, then dispensed into each well in 96-well culture plates. The wells were checked under a phase-contrast microscope, and those containing just one cell were marked and further checked daily. Holoclones from the marked wells were sub-cultured and used for further assays.

### Reagents

Rabbit anti-GLUL (ab197024) and rabbit anti-SMAD3 (phospho S423 + S425) (ab52903) antibodies were purchased from Abcam. Mouse anti-GAPDH (MA5-15738) antibody was from ThermoFisher Scientific. Rabbit anti-phospho-Akt (Ser473) (4060), mouse anti-phospho-p70S6Kinase (Thr389) (9206), rabbit anti-phospho-4EBP1 (Thr37/46) (2855), rabbit anti-phospho-p44/42 MAPK (ERK1/2 Thr202/Tyr204) (4370), rabbit anti-phospho-CaMK2 (Thr286) (12716), rabbit anti-phospho-SMAD2 (Ser465/467) (3108), rabbit anti-SMAD2/3 (8685), rabbit anti-SMAD4 (9515, 38454), and rabbit anti-Histone H3 (4499) were from Cell Signaling. Goat anti-CaMKIIγ (sc-1541) antibody was from Santa Cruz Biotechnology. Methionine sulfoximine (M5379), MG132 (M8699), and chloroquine (C6628) were from Sigma-Aldrich. KN-92 (9000890) and KN-93 (13319) were from Cayman Chemical. PI/RNase staining solution (4087) was from Cell Signaling. APC Annexin V apoptosis detection kit with 7-AAD (640930) was from BioLegend. TF activation profiling plate array 1 was from Signosis, Inc. Transcriptor high fidelity cDNA synthesis kit was from Roche (05091284001).

### Plasmid construction and transfection

shRNA constructs targeting human GLUL (TRCN0000045628 and TRCN0000343992) and SMAD3 (TRCN0000020011) were purchased from MISSION shRNA at Sigma-Aldrich. shRNA constructs targeting human SMAD2 and SMAD4 (Addgene plasmid #37051, 37046), as well as pBabe-puro-SMAD-Flag (Addgene plasmid #37041) were gifts from Sam Thiagalingam. Full-length cDNA of human CaMK2G was cloned into pLX304 vector (a gift from David Root (Addgene plasmid #25890)) at EcoR1/Not1 sites. pBABE-puro was a gift from Hartmut Land, Jay Morgenstern, and Bob Weinberg (Addgene plasmid #1764). The new construct is named as pBabe-puro-SMAD4 K159R-Flag. pMDLg/pRRE, pMD2.G, and pRSV-Rev were gifts from Didier Trono (Addgene plasmid #12251, 12259, and 12253). Lentivirus packaging plasmids and core shRNA or overexpression plasmid were transfected into 293T cells. Retroviral expression plasmids, pBabe-puro control, and SMAD4-flag were transfected into phoenix ampho cells by X-trmeGENE HP DNA transfection reagents (Roche 06366236001). Virus-containing cell culture medium was harvested and filtered with a 0.45 µm filter (Millipore SLHA033SS), then used to infect cells. Stable cells were selected by selective drugs and used for further experiments.

### Immunohistochemistry (IHC)

High-density lung cancer tissue microarray (TMA) was purchased from US Biomax (LCC6161). TMA slides were deparaffinized and hydrated, followed by antigen retrieval with Tris/EDTA buffer pH 9.0. The primary rabbit anti-GLUL (Abcam, ab197024) antibody was incubated with slides in a humidified chamber at 4°C overnight. The IHC reaction was detected with the Bond Polymer Refine Detection (Leica Biosystems, DS98000) and aminoethyl carbazole as chromogen for the visualization of the staining. Hematoxylin (Leica Biosystems) was used as counterstaining. The intensity of positive staining for GLUL was evaluated as negative, weak staining, moderate staining, and strong staining.

### Immunoblotting

Cell lysates with equal amounts of protein were separated by 4%–20% or 10% SDS-PAGE gel (Bio-Rad), then transferred to an Immobilon transfer membrane (GE Healthcare Life Sciences) and immunoblotted with specific antibodies as indicated. All the immunoblots were visualized by enhanced chemiluminescence (Bio-Rad).

### Cell number counting

Adherent cells were digested with trypsin and then suspended with phosphate-buffered saline (PBS) buffer. Suspended cells were then mixed with trypan blue dye (Bio-Rad, 145-0013) at 1:1 ration. Alive cells were directly counted with a hemocytometer.

### Colony formation assay

1,000 or 2,000 cells from indicated cell lines were seeded in 6-well plates. After 10–14 days’ culture, cells were stained with crystal violet (0.5% crystal violet dissolved in 20% methanol). For cell number quantification, the stained colonies were counted or solubilized in 1% sodium dodecyl sulfate (SDS) and the absorbance at 600 nm was taken by a microplate reader.

### Cell cycle and cell death assays

Adherent cells were digested with trypsin and then suspended with PBS buffer. For the cell cycle assay, cells were fixed with 70% ethanol (finial concentration) at 4°C overnight. Fixed cells were then washed with PBS and stained with PI/RNase staining solution (Cell Signaling, 4087). Samples were detected by BD FACS Caliber (BD Biosciences), and the cell cycle was further analyzed with ModFit LT™ software. For the cell death assay, suspended cells were centrifuged and stained with an APC Annexin V apoptosis detection kit with 7-AAD (640930) per the manufacturer’s instructions. Samples were detected by BD FACS Caliber (BD Biosciences), and cell death was further analyzed with FlowJo™ software.

### RNA isolation and real-time PCR

Total RNA was extracted from cells using the NucleoSpin® RNA Kit (Clontech, 740955) according to the manufacturer’s instructions. 1 µg RNA from each sample was used for cDNA synthesis with a Transcriptor high fidelity cDNA synthesis kit (Roche, 05091284001). Amplification was performed in a 15-μL reaction system using SYBR^®^ Advantage^®^ qPCR Premix (Clontech, 639676). All reactions were performed in triplicate in a Realplex2 system (Eppendorf). The relative gene expression level was quantified as described previously (Feng et al., 2010).

The sequence of each primer was as follows:

POLR2A:

F: 5’- GCAGGACGTAATAGAGG -3’

R: 5’- CGGACACGACCATAGA -3’

CaMK2A:

F: 5’- GGGGGAAACAAGAAGAGCGA -3’

R: 5’- TTCCGGGACCACAGGTTTTC -3’

CaMK2B:

F: 5’- AAGCATTCCAACATCGTGCG -3’

R: 5’- GGCTTGAGGTCTCTGTGGAC -3’

CaMK2G:

F: 5’- ACTAGAACGTGAGGCTCGGA -3’

R: 5’- ACCCTTGCATTTACTCGCCA -3’

SMAD4:

F: 5’- TGGCCCAGGATCAGTAGGT -3’

R: 5’- CATCAACACCAATTCCAGCA -3’

### Statistical analysis

Statistical analysis was performed using SPSS 13.0 for Windows. The data were presented as mean values ± standard deviation. Statistically significant differences were determined by Student’s t-test and one-way ANOVA, where appropriate, and defined as p < 0.05.

---

**Figure S1.**
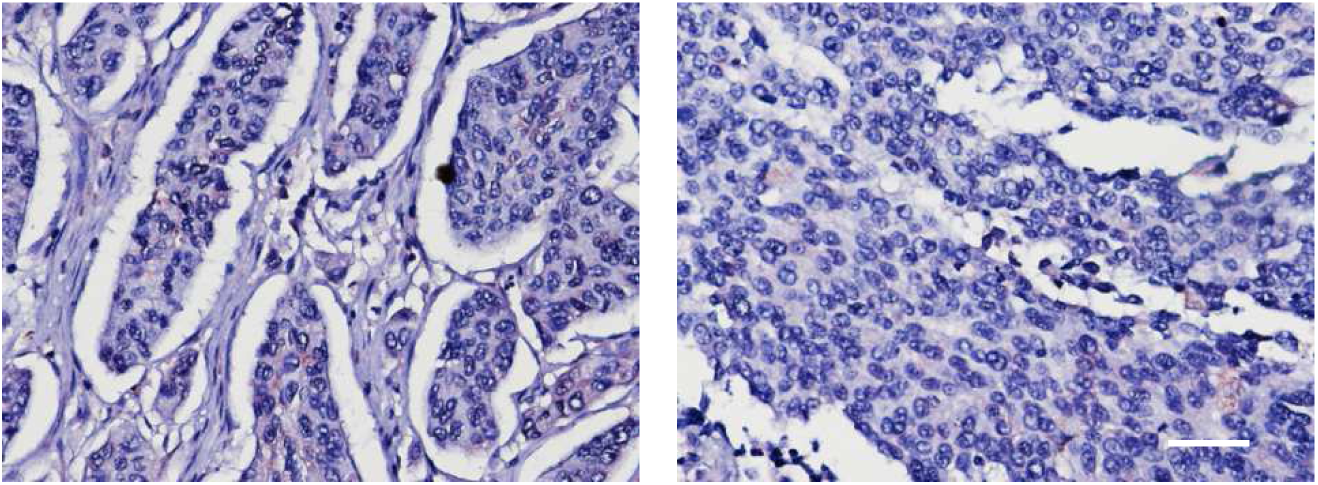
Heterogenous expression of GLUL in lung cancer samples.

**Figure S2.**
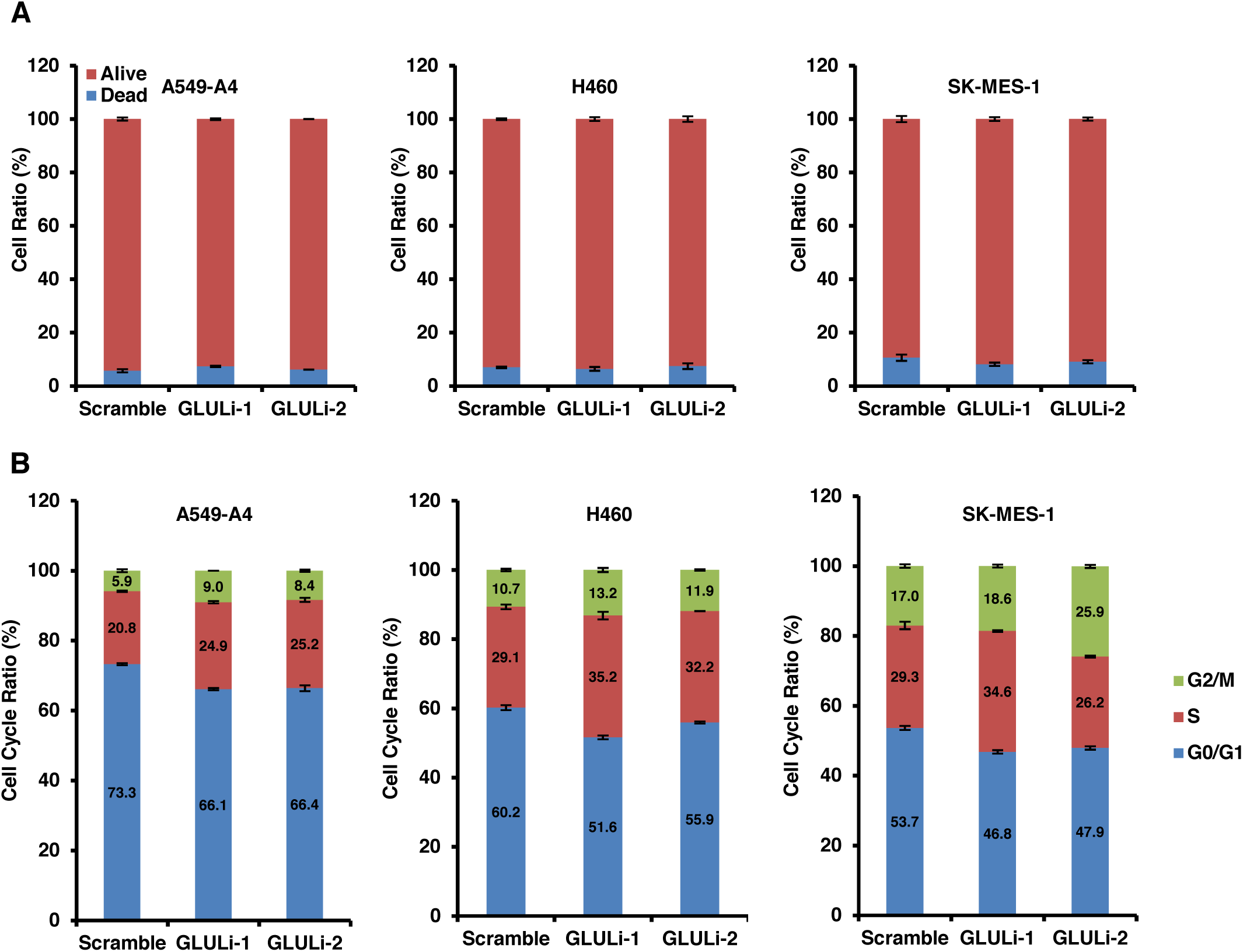
Regulation of cell death and cell cycle in GLUL-knocked-down lung cancer cells under an FM condition. (A) Cell death of GLUL-knocked-down lung cancer cells cultured with FM. (B) Cell cycle of GLUL-knocked-down lung cancer cells cultured with FM.

**Figure S3.**
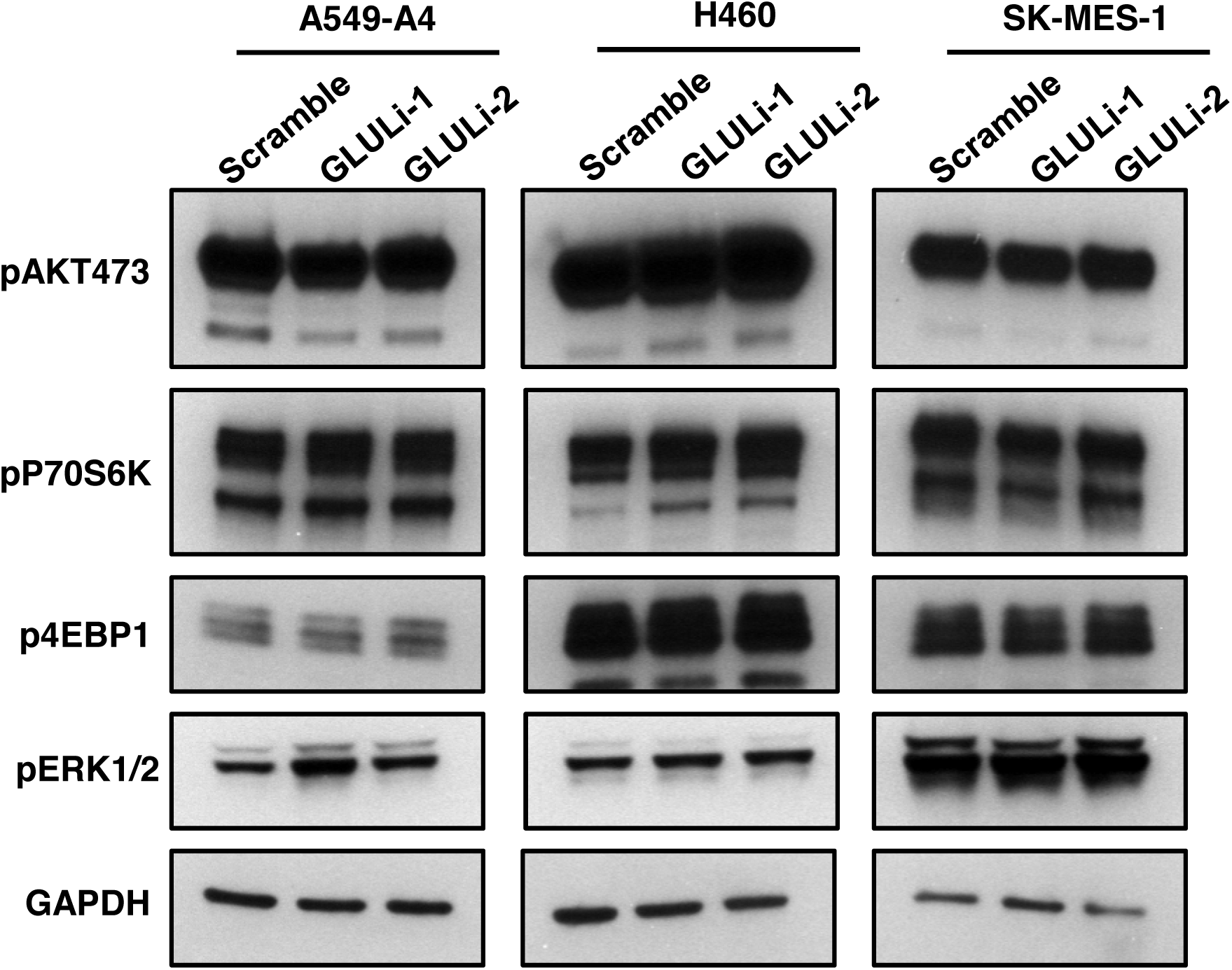
Phosphorylated key components in AKT-mTOR and ERK pathways in GLUL-knocked-down lung cancer cells.

